# High-throughput targeted gene deletion in the model mushroom Schizophyllum commune using pre-assembled Cas9 ribonucleoproteins

**DOI:** 10.1101/563759

**Authors:** Peter Jan Vonk, Natalia Escobar, Han A. B. Wösten, Luis G. Lugones, Robin A. Ohm

**Affiliations:** Microbiology, Department of Biology, Faculty of Science, Utrecht University, Utrecht, The Netherlands Padualaan 8, 3584 CH Utrecht, The Netherlands

## Abstract

Efficient gene deletion methods are essential for the high-throughput study of gene function. Compared to most ascomycete model systems, gene deletion is more laborious in mushroom-forming basidiomycetes due to the relatively low incidence of homologous recombination (HR) and relatively high incidence of non-homologous end-joining (NHEJ). Here, we describe the use of pre-assembled Cas9-sgRNA ribonucleoproteins (RNPs) to efficiently delete the homeodomain transcription factor gene hom2 in the mushroom-forming basidiomycete Schizophyllum commune by replacing it with a selectable marker. All components (Cas9 protein, sgRNA, and repair template with selectable marker) were supplied to wild type protoplasts by PEG-mediated transformation, abolishing the need to optimize the expression of cas9 and sgRNAs. A Δku80 background further increased the efficiency of gene deletion. A repair template with homology arms of 250 bp was sufficient to induce homologous recombination, whereas 100 bp was not. This is the first report of the use of pre-assembled Cas9 RNPs in a mushroom-forming basidiomycete and this approach may also improve the genetic accessibility of non-model species.

## Introduction

Mushrooms are a nutritious food source that can be cultivated on waste streams, making them important for sustainable food production^1^. Furthermore, they can produce pharmacological compounds, lignocellulolytic enzymes and show potential as cell factories^2^. Their total economic value is estimated at 63 billion USD^3^. However, the biological mechanisms underlying mushroom formation are poorly understood, primarily due to a lack of efficient genetic tools, which are essential for a deep understanding of these developmental processes^4–9^. Next-generation sequencing has yielded a vast amount of genome and expression data, but there is a relative lack of functional characterization of gene function in mushroom-forming fungi^10–14^.

The basidiomycete *Schizophyllum commune* is a model organism for mushroom formation: it completes its lifecycle in the lab in ten days and many genomic and genetic tools are available ^4,10,15^. Targeted gene deletion succeeded in four mushroom-forming species to date ^5,8,16,17^ and the majority were made in *S. commune*, although even in this organism only 23 gene deletions have been reported ^4–6,15,18–26^. The main bottleneck in gene deletion in mushroom-forming fungi is the relatively low incidence of homologous recombination (HR) and relatively high incidence of non-homologous end-joining (NHEJ)^5^. This results in the frequent ectopic integration of plasmids, instead of the desired targeted gene deletion, necessitating the laborious screening of hundreds of transformants to identify a correct gene deletion. This procedure was improved by the development of a *Δku80* strain in which the incidence of NHEJ is considerably reduced^15^, resulting in fewer transformants with an ectopic integration (and therefore fewer colonies to screen). However, the incidence of HR is not markedly increased, which means that the absolute number of transformants with a correct gene deletion does not increase. Therefore, gene deletion in mushroom-forming fungi remains a labor-intensive process. More efficient and high-throughput methods for gene deletion are required to keep up with the genomic data produced.

CRISPR-Cas9 is a powerful genetic tool greatly enhancing the efficiency of genome editing in a large number of species^27^. This endonuclease can be programmed by a ~100 bp single guide RNA (sgRNA) to target DNA based on a 20 bp homology sequence, enhancing recombination rates to facilitate insertions, deletions and replacement at the targeted site^27^. Currently, the system is primarily used with a plasmid encoding the 4 kb *cas9* gene, an endogenous promoter, a nuclear localization signal and an endogenous terminator^28^. The sgRNA can either be expressed using a RNA Polymerase III promoter or transcribed *in vitro* and co-transformed into the target cell ^27, 29^. This method has been applied to several fungi, including the mushrooms *Coprinopsis cinerea* and *Ganoderma lucidum*, where reporter genes have been deleted^17,28^. However, this procedure requires extensive genetic knowledge of the target species for codon-optimization, expression and sgRNA delivery. Moreover, methods that allow the efficient deletion of any gene (as opposed to a counter-selectable reporter gene) have not been reported in multicellular basidiomycetes.

An alternative to expressing the *cas9* gene and the sgRNA is to use pre-assembled ribonucleoprotein (RNP) complexes of Cas9 protein and *in vitro* transcribed sgRNAs^30,31^. In fungi this method has been applied to several yeasts and multicellular ascomycetes, including multiple *Aspergillus* species, *Penicillium chrysogenum* and *Cryptococcus neoformans*^32–36^.

Here, we describe the use of pre-assembled Cas9-sgRNA RNPs to delete the homeodomain transcription factor *hom2* in the basidiomycete *S. commune* with an unprecedented efficiency for mushroom-forming fungi. Although a *Δku80* background increases the efficiency of gene deletion, it is not required, which improves the genetic accessibility of non-model species. To our knowledge, this is the first report of the use of pre-assembled Cas9-sgRNA RNPs in a mushroom-forming basidiomycete.

## Results

### Cas9 RNPs are not sufficient for efficient gene disruption without a positive selection marker

The minimal form of gene inactivation using CRISPR-Cas9 is the introduction of errors by NHEJ-mediated repair after the introduction of a double-stranded break^37^. We attempted this approach by disrupting a previously introduced nourseothricin resistance cassette in the genome of strain H4.8A *Δhom2^6^.* Two sgRNAs targeting the first 100bp of the nourseothricin resistance gene were designed and transcribed *in vitro.* The efficacy of the sgRNAs was confirmed by *in vitro* digestion of the nourseothricin gene. The sgRNAs were assembled with Cas9 and introduced in H4.8A *Δhom2* protoplasts using PEG-mediated transformation. After regeneration, the protoplasts were plated on non-selective medium. The resulting colonies were tested for a loss of nourseothricin resistance, which would indicate a disruption of the nourseothricin resistance cassette by the RNPs. However, despite repeated attempts no nourseothricin sensitive colonies were identified among over 100 potential mutants, indicating a lack of gene-inactivating mutations. This means that the efficiency of the procedure is less than 1%, which strongly reduces its use in high-throughput gene deletion.

### Co-transformation of Cas9 RNPs with a repair template greatly increases the rate of gene inactivation

Next, we determined whether Cas9-sgRNA RNPs were able to increase the rate of gene deletions, when used in combination with a repair template containing a selection marker. We used the previously published *hom2* deletion vector, which is comprised of a nourseothricin antibiotic resistance cassette flanked by approximately 1200 bp flanks of the hom2 locus (Figure 1). Furthermore, it contains a phleomycin antibiotic resistance cassette, allowing us to distinguish between the desired gene deletion by homologous recombination (nourseothricin resistant and phleomycin sensitive transformants) and the undesired ectopic integration of the full construct (nourseothricin and phleomycin resistant transformants). This approach was used previously to generate a single deletion strain of the *hom2* gene that grows faster and is not capable of forming mushrooms as a dikaryon^6^. Two sgRNAs were designed that target the *hom2* gene 36 and 57 bp downstream of the translation start site, introducing double-stranded DNA breaks (Figure 1). The efficacy of the sgRNAs was confirmed by *in vitro* digestion of the *hom2* locus. Wild type H4-8 protoplasts were transformed with the *hom2* deletion vector, with or without Cas9-sgRNA RNP complexes. After regeneration, the protoplasts were plated on medium containing nourseothricin and the resulting colonies were screened for phleomycin resistance. Similar numbers of nourseothricin resistant and phleomycin sensitive transformants were found (Table 2). However, seven *hom2* gene deletions were observed with Cas9 RNPs as determined by PCR, while no gene deletions were observed without Cas9 in three replicate experiments (Table 2). Sequencing the genomic locus of these potential gene deletion mutants indeed showed replacement of the *hom2* gene with the nourseothricin resistance cassette in all cases. This indicates that the doublestranded DNA breaks introduced by the Cas9 RNPs increase the rate of gene deletion by homologous recombination.

**Table 2.** Summary of results across three replicates per condition.

| **Strain** | **Cas9**<br>**RNP** | **Nourseothricin**<br>**resistant**<br>**transformants** | **Nourseothricin**<br>**resistant and**<br>**phleomycin**<br>**sensitive**<br>**transformants** | **Confirmed gene**<br>**deletions** |
| --- | --- | --- | --- | --- |
| ***H4-8A*** | No | 103 | 24 | 0 |
| ***H4-8A*** | Yes | 125 | 26 | 7 |
| ***H4-8A Δku80*** | No | 4 | 1 | 0 |
| ***H4-8A Δku80*** | Yes | 120 | 60 | 30 |

**Figure 1.**
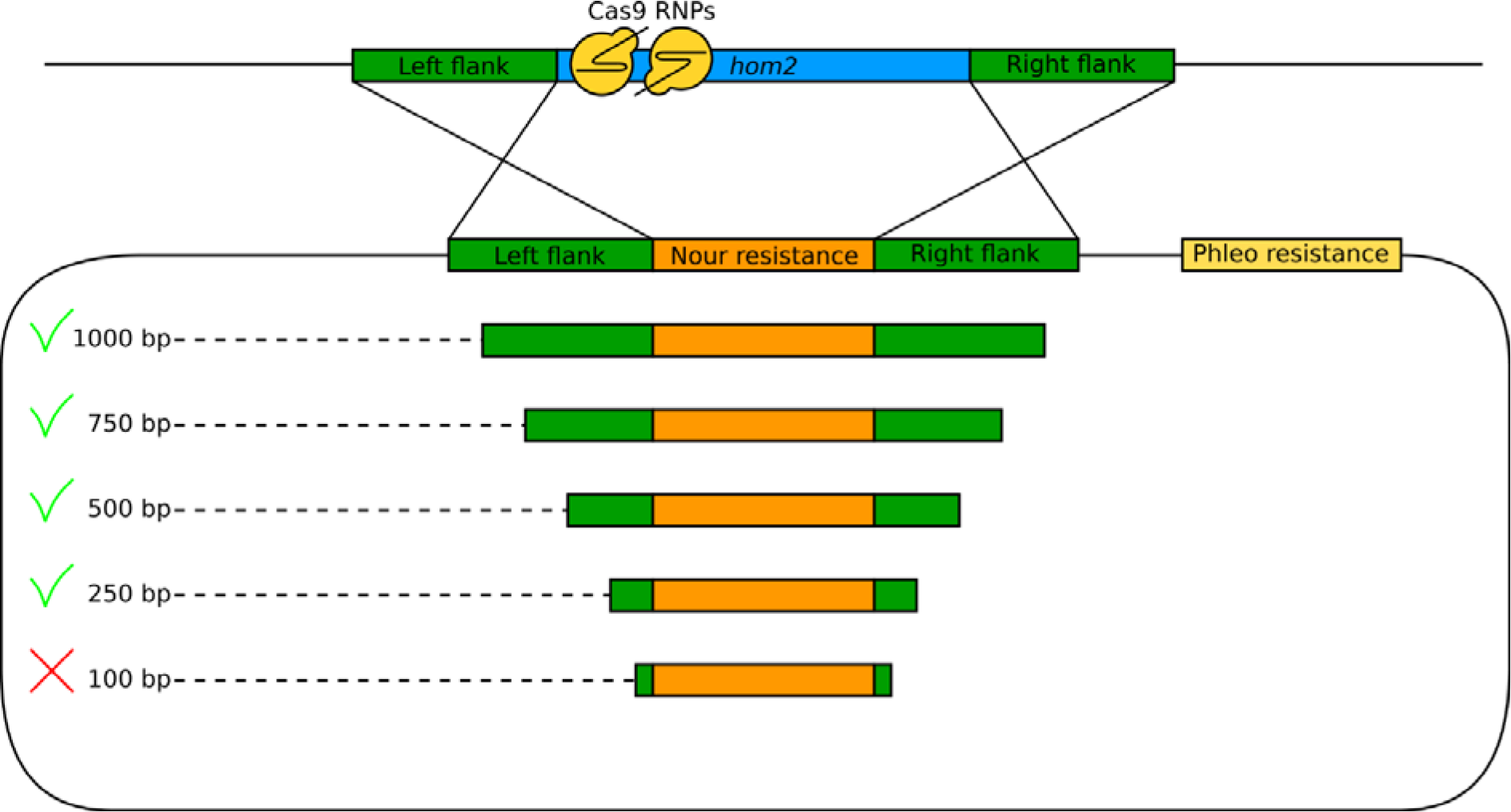
Schematic overview of the approach for targeted deletion of the *hom2* gene. Two single guide RNAs (sgRNAs) were designed that target the 5’ region of the *hom2* gene. A circular deletion vector was used as a repair template, comprised of homology arms of approximately 1200 bp flanking a nourseothricin resistance cassette. It also contains a phleomycin resistance cassette that is used to distinguish between the desired gene deletion by homologous recombination (nourseothricin resistant and phleomycin sensitive transformants) and the undesired ectopic integration of the full construct (nourseothricin and phleomycin resistant transformants). Moreover, linear PCR products with shorter homology arms were used as repair template, as indicated. The Cas9 protein, sgRNAs and repair template were introduced into protoplasts by PEG-mediated transformation. All repair templates resulted in confirmed gene deletions, with the exception of the template with 100 bp homology arms.

### A *Δku80* background further improves gene deletion efficiency

Current gene deletion methods in Basidiomycota often rely on a *Δku80* background, which inactivates the NHEJ pathway and greatly reduces the number of transformants with ectopic integrations. Therefore, we attempted the same transformation method in S. *commune Δku80.* As expected, few nourseothricin resistant transformants were observed without the addition of Cas9 RNPs and no confirmed gene deletions were identified among these (Table 2). In the *Δku80* strain co-transformed with Cas9 RNPs, a similar number of nourseothricin resistant transformants was observed as in the wild type strain with Cas9 (Table 2). However, both the number of phleomycin sensitive transformants and the number of transformants with a confirmed gene deletion were significantly higher (Table 2), with a total of 30 confirmed gene deletions. Sequencing of the genomic locus of 10 *hom2* deletion strains confirmed the expected deletion of the *hom2* gene and insertion of the nourseothricin resistance cassette in all cases. This indicates that using Cas9 RNPs in a *Δku80* background increases the rate of homologous recombination and results in a higher rate of gene deletion than in the wild type.

### Homology arms of 250 bp are sufficient for the recruitment of homology-directed repair

Current gene deletion methods in mushroom-forming Basidiomycota generally rely on repair templates comprised of homology arms of approximately 1000 bp that flank a selectable marker. Indeed, the *hom2* deletion vector described above contains homology arms of 1190 and 1147 bp. We determined whether repair templates with shorter homology arms can also assist in gene deletion when used in combination with Cas9-sgRNA RNPs. To generate the repair templates, PCRs were performed on the *hom2* deletion vector resulting in linear double-stranded DNA fragments with reduced homology arm lengths of approximately 1000, 750, 500, 250 and 100 bp (Figure 1). For all homology arm lengths at least 2 confirmed gene deletions were obtained, except for 100 bp homology, for which no gene deletions were observed despite repeated attempts. This indicates that 250 bp, but not 100 bp, homology is sufficient to recruit homology-directed repair, resulting in a gene deletion.

## Discussion

Efficient gene deletion methods are essential for the high-throughput study of gene function. Here, we described a method based on a pre-assembled ribonucleoprotein (RNP) complex of Cas9 protein and *in vitro* transcribed short guide RNAs. This method was used to create 47 deletion strains of the *hom2* gene by replacing it with a selectable marker. Moreover, we determined that homology arms of 250 bp are sufficient to efficiently induce homologous recombination. This method yields considerably more gene deletion strains than our previously published methods^5,15^, in which usually only 1 correct deletion strain was obtained and only after repeated attempts. Various efficient approaches of genome editing using CRISPR/Cas9 have been reported in the model fungus *Saccharomyces cerevisiae.* For example, *cas9* and sgRNAs may be expressed from a plasmid and cause genome edits with such high efficiency that the use of a selectable marker is not necessary^38^. We have attempted this approach in *S. commune*, but were unsuccessful in producing strains with a targeted gene deletion (data not shown). Nor were we successful with *in vitro* produced Cas9 RNPs without the use of a selectable marker, since no strains with a gene deletion were identified among > 100 colonies. This suggests that the efficiency of the Cas9 RNP approach is much lower in *S. commune* (and possibly other mushroom-forming basidiomycetes) than in Ascomycota. This relatively low efficiency is also observed in *Coprinopsis cinerea^28^.* In contrast, many correct gene deletion strains were obtained when we attempted to replace the target gene with a selectable marker. We hypothesize that the Cas9 RNPs cause double strand breaks in the target gene, after which the DNA repair machinery is recruited to the site and repairs the damage using the provided repair template. Although this is a relatively rare event, positive selection subsequently allows the identification of the (relatively small number of) correct transformants.

Current gene deletion methods in mushroom-forming Basidiomycota generally rely on templates comprised of homology arms of approximately 1000 bp that flank a selectable marker^5,16^. The creation of this gene deletion template is currently a bottleneck in high throughput gene deletion, although recent approaches for the assembly of multiple DNA fragments (e.g. Gibson assembly) have considerably improved this^39^. Our results show that homology arms as short as 250 bp are effective in inducing homologous recombination when used in combination with RNPs. This relatively short length of the template is well within the range of cost-effective chemical synthesis, which will further facilitate high-throughput gene deletion by allowing *in vitro* construction of this template.

The methods described in this study will remove several bottlenecks for efficient genome editing in non-model mushroom-forming fungi, as well as wild isolate strains of *S. commune.* Firstly, our approach works efficiently not only in a *Δku80* background, but also in the wild type. This will simplify the gene deletion procedure for non-model system species for which no *Δku80* strains are available. Secondly, the expression levels of *cas9* and the sgRNAs do not need to be optimized using organism-specific promoters, since both components are supplied externally. Previous studies have shown that the selection markers needed for the repair template are functional in other mushroom-forming fungi^40^.

In addition to gene deletion, these techniques can be modified to produce other genome edits, including promoter replacement and targeted gene integration (knock-in). Recent efforts in genome and transcriptome sequencing of mushroom-forming species have resulted in a large number of candidate gene families involved in mushroom development and other processes ^11,14,41,42^. The important addition of Cas9 RNP-mediated genome editing to the molecular toolkit of *S. commune* will facilitate the functional characterization of these gene families, leading to important new insights into the biology of this important group of fungi.

## Materials & methods

### Culture conditions and strains

*S. commune* was grown from a small agar inoculum in *Schizophyllum commune* minimal medium (SCMM) at 30 °C^43^. For solid culture, the medium was supplemented with 1.5% agar. All strains are derived from *S. commune* H4-8 (matA43matB41; FGSC 9210)^5,10^. Strain *Δku80*^15^ and strain *Δhom2*^6^ were published previously. For selection on nourseothricin (Bio-Connect, Netherlands) or phleomycin (Bio-Connect, Netherlands), SCMM was supplemented with 15 μg ml^−1^ and 25 μg ml^−1^ antibiotic, respectively^40^.

### Cas9 expression and purification

pET-NLS-Cas9-6xHis was a gift from David Liu (Addgene plasmid # 62934; http://n2t.net/addgenei62934; RRID:Addgene_62934)^44^ and contains a NLS-Cas9-6xhis fusion sequence under control of the *17* promotor. The plasmid was introduced into *Escherichia coli* BL21 Star (DE3) (Life Technologies, USA). The resulting expression strain was pre-grown for 16 hours in liquid culture at 37 °C in LB supplemented with 50 μg ml^−1^ ampicillin. This pre-culture was diluted 1:100 in fresh LB with ampicillin and grown in liquid culture at 22.5 °C to an OD_600_ of 0.6 when Cas9 expression was induced with IPTG at a final concentration of 0.4 mM and incubation was continued for an additional 18 hours. Cells were harvested by centrifugation and resuspended in 80 ml wash buffer (20 mM Tris-HCl pH 8.0, 30 mM imidazole, 500 mM NaCl) supplemented with 1 mM PMFS, 1 mM MgCl_2_ and 6.25 U ml^−1^ DNase I (ThermoFisher Scientific, USA) and lysed three times with a French press (AMINCO, USA) at 500 psi on high setting in a pre-chilled cylinder (4 °C). After lysis, all steps were performed at 4 °C. The lysate was cleared of cellular debris by centrifugation for 30 minutes at 15,000 g. Per liter of culture, 8 ml of bed volume of Ni-NTA resin (Qiagen, Germany) was added and incubated for one hour with gentle agitation. Column purification was performed according to the manufacturer’s recommendations and Cas9 was eluted in 24 ml wash buffer with 500 mM imidazole. The protein purity was assessed on SDS-PAGE with colloidal coomassie staining^45^ and protein yield was quantified by BCA-assay (ThermoFisher Scientific, USA). Imidazole and salts were removed by three washing steps with 21 ml dialysis buffer (20 mM Tris-HCl pH 8.0, 200 mM KCl, 10 mM MgCl_2_) on Amicon 30 kDa filters (Merck Millipore, USA). Cas9 protein was diluted to 10 mg ml^−1^ in dialysis buffer, frozen in liquid nitrogen and stored at −80 °C until use.

### Design and synthesis of gene-specific sgRNAs for gene disruption

Candidate protospacers (including PAM site) were identified in the coding region of the nourseothricin resistance cassette^43^ and *hom2* (protein ID 257987 in version Schco3 of the genome of *S. commune*^10^) with CCtop^46^ and checked against the full genome to identify potential off-target regions (defined here as regions with fewer than 5 bp differences with the actual target region as identified by Bowtie2^47^). Two sgRNAs were selected per gene based on fewest off-targets and the presence of one or more guanines at the start of the sgRNA as this promotes a high yield of *in vitro 17* transcription. For the nourseothricin resistance gene, sgRNA 1 and 2 were located 57 and 22 bp downstream of the translation start site, respectively, and for *hom2*, sgRNA 1 and 2 were located 57 and 36 bp downstream of the translation start site of *hom2*, respectively (Figure 1). Oligos were designed (p1_sgRNA and p2_sgRNA; Table 1) and the sgRNAs were synthesized *in vitro* according to the specifications of the GeneArt Precision sgRNA Synthesis Kit (ThermoFisher Scientific, USA).

**Table 1.**
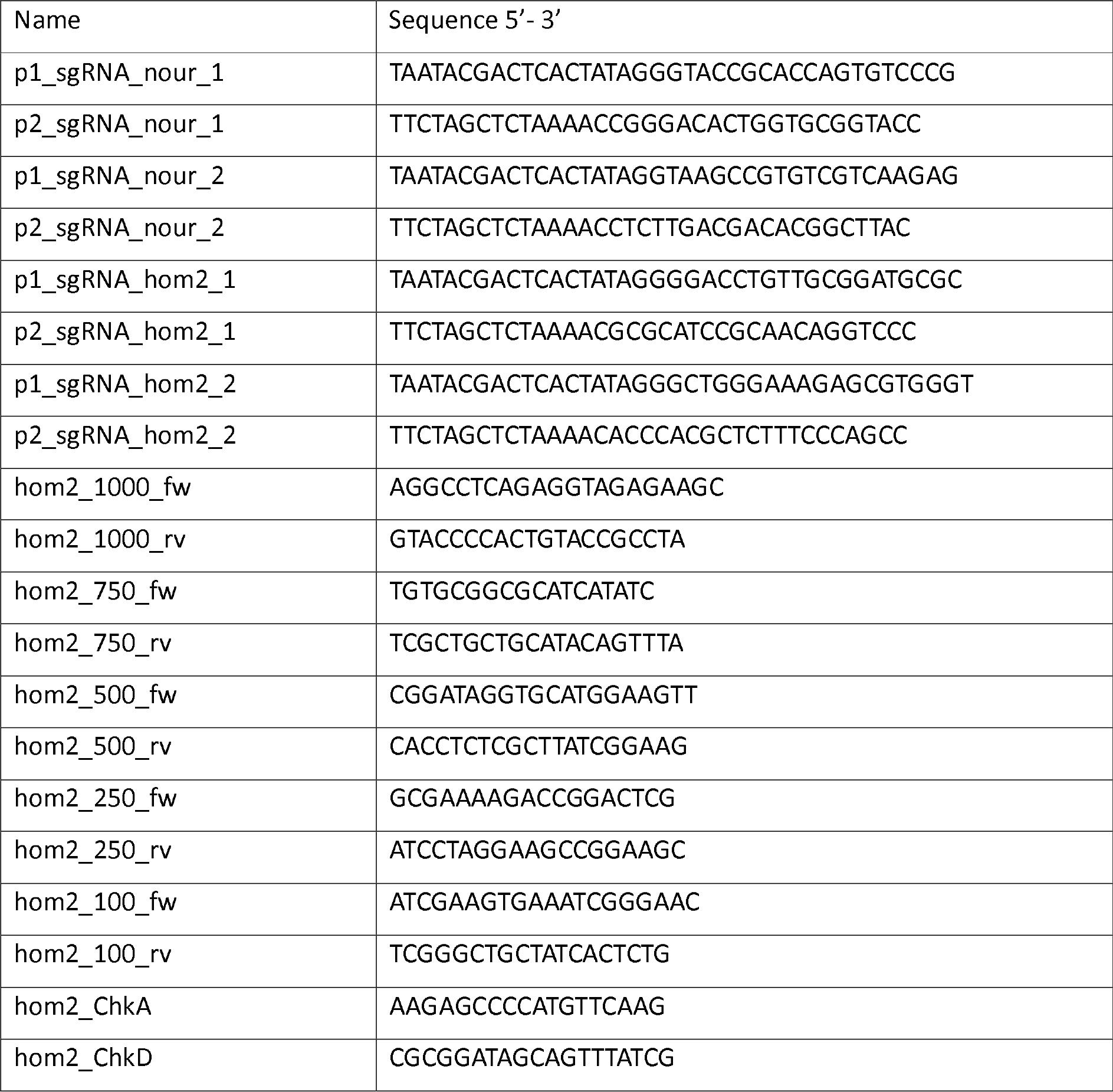
Primers used in this study

### *In vitro* Cas9 digestion assay

The efficacy of the RNPs was tested by an *in vitro* digestion of a template containing the target regions. For the nourseothricin resistance cassette, the resistance gene was cut from the pDELCAS deletion vector^5^ with EcoRI, resulting in a 3759 bp fragment. *In vitro* digestion by the Cas9 sgRNA RNPs results in two fragments of 2719 bp and 1040 bp for sgRNA 1 and 2684 bp and 1075 bp for sgRNA 2. The template for *hom2* was generated by PCR with primers annealing to the promotor and terminator region of *hom2* (hom2_250_fw and hom2_250_rv). This resulted in a 3656 bp fragment that can be digested by Cas9 with either sgRNA to two fragments of 433 bp and 3223 bp for sgRNA 1 and 412 bp and 3244 bp for sgRNA 2. For digestion, 1 μg of Cas9, 200 ng sgRNA (which is approximately a 1:1 molar ratio) and 1× Cas9 reaction buffer (20 mM HEPES pH 6.5, 100 mM NaCl, 5 mM MgCl_2_ 0.1 mM EDTA) were mixed in 25 μl final volume and assembled at 37 °C for 10 minutes. After assembly, 500 ng of template (100 ng μl^−1^) was added to a final volume of 30 μl and the sample was incubated at 37 °C for 15 minutes. Cas9 was degraded by the addition of 1 U Proteinase K (ThermoFisher Scientific, USA) and incubated at room temperature for 15 minutes, after which DNA digestion was verified on agarose gel.

### Design of the repair templates for homologous recombination

The previously described *hom2* deletion vector was used as a template for homologous recombination^6^. This plasmid contains a nourseothricin resistance cassette flanked by 1200 bp homology arms outside *hom2.* Moreover, the plasmid harbors a phleomycin resistance cassette that is only integrated if the plasmid integrates through a single cross-over (i.e. ectopically). Linear templates with reduced homology arm lengths (approximately 1000 bp, 750 bp, 500 bp, 250 bp and 100 bp) were made by PCR on the full vector. The primers were designed to bind 1000 bp (hom2_1000_fw and hom2_1000_rv; Table 1), 750 bp (hom2_750_fw and hom2_750_rv; Table 1), 500 bp (hom2_500_fw and hom2_500_rv; Table 1), 250 bp (hom2_250_fw and hom2_250_rv; Table 1) and 100 bp (hom2_100_fw, hom2_100_rv; Table 1) outside of the nourseothricin resistance cassette (Figure 1).

### Transformation of *S. commune*

Protoplasts were prepared as previously described and stored at −80 °C until use^43^. 20 μg Cas9 was mixed with the two sgRNAs (2 μg of each sgRNA) targeting the gene of interest (resulting in a 2:1:1 molar ratio of Cas9, sgRNA 1 and sgRNA 2) and 1× Cas9 buffer (20 mM HEPES, 100 mM NaCl, 5 mM MgCl_2_, 0.1 mM EDTA, pH 6.5). In the negative controls, Cas9 was replaced with dialysis buffer and sgRNA with MilliQ water. Cas9 and sgRNAs were pre-assembled for 10 minutes at 37 °C.

In the case of the nourseothricin gene inactivation, 4 × 10^3^ protoplasts in 200 μl were added. Subsequent transformation was performed as previously described, but without selection^6^. The regenerated protoplasts were plated on non-selective medium and grown for two days.

In the case of *hom2* deletions, the number of protoplasts was changed to 2 × 10^7^ in 200 μl and 12.5μg repair template was added. The regenerated protoplasts were plated on medium supplemented with nourseothricin and grown for 3 days. Next, 48 colonies were randomly selected per transformation after three days and transferred to fresh selection media for a second round of selection. In the case of transformation with a full *hom2* deletion plasmid, nourseothricin-resistant colonies were then transferred to phleomycin selection medium and their resistance was scored. DNA was isolated from agar plugs by microwaving samples for 120 seconds in 50 μl TE buffer (10 mM Tris pH 8.0, 1 mM EDTA) from candidates that were resistant to nourseothricin and sensitive to phleomycin for PCR verification with primers before the upstream homology arm (hom2_ChkA; Table 1) and after the downstream homology arm (hom2_ChkD; Table 1)^6,48^. The size of the DNA band was used to determine whether the gene of interest was replaced with the nourseothricin resistance cassette. Primers hom2_100_fw and hom2_100_rv (Table 1) were used to sequence the PCR fragment to confirm the correct integration.

## Acknowledgements

This project has received funding from the European Research Council (ERC) under the European Union’s Horizon 2020 research and innovation programme (grant agreement number 716132).

## Contributions

RO, LL and HW conceived the project. PV and NE performed experiments. PV and RO analyzed the data. PV and RO wrote the manuscript. All authors read and approved the manuscript. RO provided funding.

## Competing interests

The authors declare no competing interests.

## Data availability

The datasets generated during and/or analysed during the current study are available from the corresponding author on reasonable request.

## References

1. Grimm, D. & Wösten, H. A. B. Mushroom cultivation in the circular economy. Appl. Microbiol. Biotechnol. 102, 7795–7803 (2018).

2. de Mattos-Shipley, K. M. J. et al. The good, the bad and the tasty: The many roles of mushrooms. Stud. Mycol. 85, 125–157 (2016).

3. Royse, D. J., Baars, J. & Tan, Q. Current Overview of Mushroom Production in the World. in Edible and Medicinal Mushrooms 5–13 (John Wiley & Sons, Ltd, 2017). doi:10.1002/9781119149446.ch2

4. Pelkmans, J. F. et al. Transcription factors of Schizophyllum commune involved in mushroom formation and modulation of vegetative growth. Sci. Rep. 7, 310 (2017).

5. Ohm, R. A. et al. An efficient gene deletion procedure for the mushroom-forming basidiomycete Schizophyllum commune. World J. Microbiol. Biotechnol. 26, 1919–1923 (2010).

6. Ohm, R. A., de Jong, J. F., de Bekker, C., Wösten, H. A. B. & Lugones, L. G. Transcription factor genes of Schizophyllum commune involved in regulation of mushroom formation. Mol. Microbiol. 81, 1433–1445 (2011).

7. Collins, C. M., Heneghan, M. N., Kilaru, S., Bailey, A. M. & Foster, G. D. Improvement of the Coprinopsis cinerea molecular toolkit using new construct design and additional marker genes. J. Microbiol. Methods 82, 156–162 (2010).

8. Nakazawa, T., Ando, Y., Kitaaki, K., Nakahori, K. & Kamada, T. Efficient gene targeting in ∆Cc.ku70 or ∆Cc.lig4 mutants of the agaricomycete coprinopsis cinerea. Fungal Genet. Biol. 48, 939–946 (2011).

9. Kües, U. & Navarro-González, M. How do Agaricomycetes shape their fruiting bodies? 1. Morphological aspects of development. Fungal Biol. Rev. 29, 63–97 (2015).

10. Ohm, R. A. et al. Genome sequence of the model mushroom Schizophyllum commune. Nat. Biotechnol. 28, 957–963 (2010).

11. Grigoriev, I. V. et al. MycoCosm portal: gearing up for 1000 fungal genomes. Nucleic Acids Res. 42, D699–D704 (2014).

12. Stajich, J. E. et al. Insights into evolution of multicellular fungi from the assembled chromosomes of the mushroom Coprinopsis cinerea (Coprinus cinereus). Proc Natl Acad Sci U S A 107, 11889–11894 (2010).

13. Morin, E. et al. Genome sequence of the button mushroom Agaricus bisporus reveals mechanisms governing adaptation to a humic-rich ecological niche. Proc. Natl. Acad. Sci. 109, 17501–17506 (2012).

14. Nagy, L. G., Kovács, G. M. & Krizsán, K. Complex multicellularity in fungi: Evolutionary convergence, single origin, or both? Biological Reviews (2018). doi:10.1111/brv.12418

15. De Jong, J. F., Ohm, R. A., De Bekker, C., Wösten, H. A. B. & Lugones, L. G. Inactivation of ku80 in the mushroom-forming fungus Schizophyllum commune increases the relative incidence of homologous recombination. FEMS Microbiol. Lett. 310, 91–95 (2010).

16. Salame, T. M. et al. Predominance of a versatile-peroxidase-encoding gene, mnp4, as demonstrated by gene replacement via a gene targeting system for Pleurotus ostreatus. Appl. Environ. Microbiol. 78, 5341–5352 (2012).

17. Qin, H., Xiao, H., Zou, G., Zhou, Z. & Zhong, J. J. CRISPR-Cas9 assisted gene disruption in the higher fungus Ganoderma species. Process Biochem. 56, 57–61 (2017).

18. Schubert, D., Raudaskoski, M., Knabe, N. & Kothe, E. Ras GTPase-Activating Protein Gap1 of the Homobasidiomycete Schizophyllum commune Regulates Hyphal Growth Orientation and Sexual Development. Eukaryot. Cell 5, 683–695 (2006).

19. Van Peer, A. F. et al. The septal pore cap is an organelle that functions in vegetative growth and mushroom formation of the wood-rot fungus Schizophyllum commune. Environ. Microbiol. 12, 833–844 (2009).

20. Knabe, N. et al. A central role for Ras1 in morphogenesis of the basidiomycete Schizophyllum commune. Eukaryot. Cell 12, 941–52 (2013).

21. Robertson, C. I., Bartholomew, K. A., Novotny, C. P. & Ullrich, R. C. Deletion of the Schizophyllum commune A alpha locus: the roles of A alpha Y and Z mating-type genes. Genetics 144, 1437–44 (1996).

22. Wetter, M.-A., Schuren, F. H. J., Schuurs, T. A. & Wessels, J. G. H. Targeted mutation of the SC3 hydrophobin gene of Schizophyllum commune affects formation of aerial hyphae. FEMS Microbiol. Lett. 140, 265–269 (1996).

23. Horton, J. S., Palmer, G. E. & Smith, W. J. Regulation of Dikaryon-Expressed Genes byFRT1in the BasidiomyceteSchizophyllum commune. Fungal Genet. Biol. 26, 33–47 (1999).

24. Lengeler, K. B. & Kothe, E. Identification and characterization of brt1, a gene down-regulated during B-regulated development in Schizophyllum commune. Curr. Genet. 35, 551–6 (1999).

25. Lengeler, K. B. & Kothe, E. Mated: a putative peptide transporter of Schizophyllum commune expressed in dikaryons. Curr. Genet. 36, 159–64 (1999).

26. Lugones, L. G. et al. The SC15 protein of Schizophyllum commune mediates formation of aerial hyphae and attachment in the absence of the SC3 hydrophobin. Mol. Microbiol. 53, 707–716 (2004).

27. Jinek, M. et al. A Programmable Dual-RNA-Guided DNA Endonuclease in Adaptive Bacterial Immunity. Science (80-.). 337, 816–821 (2012).

28. Sugano, S. S. et al. Genome editing in the mushroom-forming basidiomycete Coprinopsis cinerea, optimized by a high-throughput transformation system. Sci. Rep. 7, 1260 (2017).

29. Hruscha, A. et al. Efficient CRISPR/Cas9 genome editing with low off-target effects in zebrafish. Development 140, 4982–4987 (2013).

30. Woo, J. W. et al. DNA-free genome editing in plants with preassembled CRISPR-Cas9 ribonucleoproteins. Nat. Biotechnol. 33, 1162–1164 (2015).

31. Gasiunas, G., Barrangou, R., Horvath, P. & Siksnys, V. Cas9-crRNA ribonucleoprotein complex mediates specific DNA cleavage for adaptive immunity in bacteria. Proc. Natl. Acad. Sci. 109, E2579–E2586 (2012).

32. Grahl, N., Demers, E. G., Crocker, A. W. & Hogan, D. A. Use of RNA-Protein Complexes for Genome Editing in Non- albicans Candida Species. mSphere 2, e00218–17 (2017).

33. Song, L. et al. Efficient genome editing using tRNA promoter-driven CRISPR/Cas9 gRNA in Aspergillus niger. PLOS ONE 13, (Public Library of Science, 2018).

34. Pohl, C., Kiel, J. A. K. W., Driessen, A. J. M., Bovenberg, R. A. L. & Nygård, Y. CRISPR/Cas9 Based Genome Editing of Penicillium chrysogenum. ACS Synth. Biol. 5, 754–764 (2016).

35. Pohl, C., Mózsik, L., Driessen, A. J. M., Bovenberg, R. A. L. & Nygård, Y. I. Genome editing in penicillium chrysogenum using cas9 ribonucleoprotein particles. in Methods in Molecular Biology 1772, 213–232 (Humana Press, New York, NY, 2018).

36. Wang, Y. et al. A ‘suicide’ CRISPR-Cas9 system to promote gene deletion and restoration by electroporation in Cryptococcus neoformans. Sci. Rep. 6, 31145 (2016).

37. Ran, F. A. et al. Genome engineering using the CRISPR-Cas9 system. Nat. Protoc. 8, 2281–2308 (2013).

38. Mans, R. et al. CRISPR/Cas9: a molecular Swiss army knife for simultaneous introduction of multiple genetic modifications in Saccharomyces cerevisiae. FEMS Yeast Res. 15, (2015).

39. Gibson, D. G. Enzymatic Assembly of Overlapping DNA Fragments. Methods Enzymol. 498, 349–361 (2011).

40. Alves, A. M. C. R. et al. Highly efficient production of laccase by the basidiomycete Pycnoporus cinnabarinus. Appl. Environ. Microbiol. 70, 6379–6384 (2004).

41. Krizsan, K. et al. Transcriptomic atlas of mushroom development highlights an independent origin of complex multicellularity. bioRxiv 349894 (2018). doi:10.1101/349894

42. Vonk, P. J. & Ohm, R. A. The role of homeodomain transcription factors in fungal development. Fungal Biol. Rev. 32, 219–230 (2018).

43. Van Peer, A. F., De Bekker, C., Vinck, A., Wösten, H. A. B. & Lugones, L. G. Phleomycin increases transformation efficiency and promotes single integrations in schizophyllum commune. Appl. Environ. Microbiol. 75, 1243–1247 (2009).

44. Zuris, J. A. et al. Cationic lipid-mediated delivery of proteins enables efficient protein-based genome editing in vitro and in vivo. Nat. Biotechnol. 33, 73–80 (2015).

45. Dyballa, N. & Metzger, S. Fast and sensitive colloidal coomassie G-250 staining for proteins in polyacrylamide gels. J. Vis. Exp. (2009). doi:10.3791/1431

46. Stemmer, M., Thumberger, T., del Sol Keyer, M., Wittbrodt, J. & Mateo, J. L. CCTop: An Intuitive, Flexible and Reliable CRISPR/Cas9 Target Prediction Tool. PLoS One 10, e0124633 (2015).

47. Langmead, B. & Salzberg, S. L. Fast gapped-read alignment with Bowtie 2. Nat. Methods 9, 357–359 (2012).

48. Izumitsu, K. et al. Rapid and simple preparation of mushroom DNA directly from colonies and fruiting bodies for PCR. Mycoscience 53, 396–401 (2012).

